# MEK inhibition causes Bim stabilization and sensitivity to Bcl2 family member inhibitors in RAS-MAPK mutated neuroblastoma

**DOI:** 10.1101/2020.02.12.945089

**Authors:** Thomas F Eleveld, Lindy Vernooij, Linda Schild, Bianca Koopmans, Lindy K Alles, Marli E Ebus, Peter van Sluis, Huib N Caron, Jan Koster, Rogier Versteeg, M. Emmy M. Dolman, Jan J Molenaar

**Affiliations:** Department of Translational Research, Princess Maxima Centre for Childhood Oncology, Utrecht, the Netherlands; Department of Oncogenomics, Academic Medical Center of the University of Amsterdam, Amsterdam, the Netherlands; Department of Pediatric Oncology, Emma Children’s Hospital, Academic Medical Center of the University of Amsterdam, Amsterdam, the Netherlands

## Abstract

Mutations affecting the RAS-MAPK pathway occur frequently in relapsed neuroblastoma tumors and are associated with response to MEK inhibition *in vitro*. However, these inhibitors alone do not lead to tumor regression *in vivo*, indicating the need for combination therapy. Through high throughput combination screening we identify Trametinib and inhibitors of the BCL2 family (Navitoclax and Venetoclax) as a promising combination in neuroblastoma cells with RAS-MAPK mutations. In these lines, inhibiting the RAS-MAPK pathway leads to Bim stabilization and increased sensitivity to compounds inhibiting Bim binding to Bcl2 family members. Combining Trametinib with BCL2 inhibitors causes increased growth inhibition compared to Trametinib only in *NRAS* mutant SKNAS xenografts, while BCL2 inihibitors alone do not affect growth of these tumors. These results show that MEK inhibitors and specific Bcl2 family member inhibitors are a potent combination for RAS-MAPK mutated neuroblastoma tumors.

## Introduction

Neuroblastoma is a pediatric tumor with a highly variable prognosis: low risk neuroblastoma tumors often regress without treatment while high risk neuroblastoma tumors do not respond to intensive regimens ^1, 2^. Sequencing studies have revealed few recurrent events, and therefore little therapeutic opportunities^3-6^. Neuroblastoma relapse tumors arise from primary tumors via clonal evolution and contain more aberrations than primary tumors^7, 8^. These relapse tumors also shown an increase in targetable mutations, with the p53-MDM2-p14 ARF pathway, the *ALK* gene, and other RAS-MAPK pathway genes being frequently affected^8-11^.

Previously, we have shown that neuroblastoma cell lines containing RAS-MAPK mutations are sensitive to MEK inhibitors^*8*^. However, when used *in vitro* these inhibitors reach a plateau phase where increasing the concentration does not further affect cell viability. *In vivo* this translates to only a slight tumor reduction. Furthermore, clinical trials have shown that MEK inhibitors as a monotherapy do not lead to sustainable responses^12, 13^, warranting the search for an effective combination therapy.

Here we identify through high throughput compound screening that MEK inhibitors and Bcl2 family inhibitors are a potent combination in RAS-MAPK mutated neuroblastoma. Inhibition of the RAS-MAPK pathway causes stabilization of Bim, which is associated with increased sensitivity to Bcl2 family inhibitors. Treatment of SKNAS xenografts with the combination of Trametinib and Bcl2 inhibitors causes increased growth inhibition compared to Trametinib alone, showing that this combination is also effective *in vivo.*

## Materials and methods

### High throughput screens

Cells were seeded in 384 well plates as shown in table 1 at t=0. At t=24h Trametinib or DMSO was added to a final concentration of 1 μM. Library compounds were added at final concentrations of 10 nM, 100 nM and 1μM with appropriate solvent controls using the Sciclone ALH 3000 liquid handling robot. At t=96 hrs cell viability was determined using MTT assay. Area Under the Curve was calculated using Graphpad Prism 6.

**Table 1:**
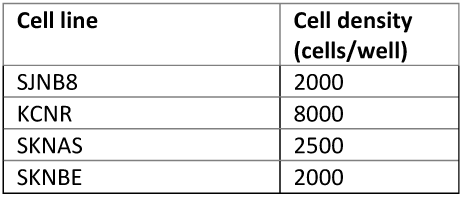
number of cells per well used for screening

| Cell line | Cell density (cells/well) |
| --- | --- |
| SJNB8 | 2000 |
| KCNR | 8000 |
| SKNAS | 2500 |
| SKNBE | 2000 |

The Sequoia anti-neoplastic library contains 157 drugs used in cancer treatment and was purchased from Sequoia Research Products. The SCREEN-WELL^®^ epigenetics library contains 43 epigenetic compounds and was acquired from Enzo Life Sciences. Other targeted inhibitors used in this study were purchased from Selleck Chemicals.

### Immunoprecipitation and Western blotting

Samples were harvested in CHAPS buffer and subjected to co-immunoprecipitation (co-IP) using Dynabeads magnetic protein A agarose beads (Life Technologies) according to the manufacturers protocol. 1 mg of protein lysate was used as input material for IP reactions. Antibodies were acquired from Cell Signaling unless noted otherwise. Antibodies were used for MEK (#8727), ERK (#9102), pMEK (#9154), pERK (#4376), Bcl2 (#4223), Bcl-XL (#2764), Mcl1 (#4572), Bim (#2933), α-tubulin (#3873) and β-actin (Abcam, Ab6276).

### Cell culture

Cell lines were cultured in DMEM supplemented with 10% FCS, 20 mM L-glutamine, 10 U/ml penicillin and 10 μg/ml streptomycin and maintained at 37 °C under 5% CO_2_. Cell line identities were regularly confirmed by short tandem repeat (STR) profiling using the PowerPlex16 system and GeneMapper software (Promega). Cell lines were regularly screened for mycoplasma.

### Cell viability assays

For cell viability assays, 5–20 × 10^3^ cells were seeded in 50 μl in 96-well plates 1 day before treatment. Trametinib to a final concentration of 1uM or DMSO was added to the cells and subsequently the Bcl2 family inhibitors were added in a seven-point fivefold dilution series. Cell viability was assayed by MTT assay after 72 hrs^8^. All experiments were performed in triplicate and values were compared to solvent-treated controls.

### Animal experiments

All experiments were conducted after obtaining ethical approval from the DEC (animal experiments committee) of the AMC under number DAG217. The SKNAS neuroblastoma cell line was transplanted subcutaneously into female NMRI nu-/nu- mice at 5–7 weeks of age. Once the engrafted tumors reached 200 mm3, mice were treated with Trametinib at 1 mg/kg in combination with 100 mg/kg of Navitoclax or Venetoclax via oral gavage. Mice were treated daily and tumor size was monitored twice a week. Tumor burden was determined according to the formula (π/6)d^3^, where d represents the mean tumor diameter obtained by caliper measurement. Statistical analysis was performed using a two-tailed t test at each time point, with P values <0.05 indicating significance between the vehicle-treated group and each of the treatment groups.

## Results

### High throughput screening of RAS-MAPK mutated cell lines in the presence and absence of Trametinib

To identify compounds that are effective in combination with MEK inhibitors in neuroblastoma, we performed a large scale drug screen. Four neuroblastoma cell lines were selected based on the presence of the most common mutations affecting the RAS-MAPK pathway, RAS (*NRAS* in SKNAS and *KRAS* in SJNB8), *NF1* (SKNBE) and *ALK* (KCNR). In total 210 compounds were screened: 60 chemotherapeutic compounds, 106 targeted drugs that are either approved for clinical use or far along in clinical development, and 44 epigenetic modifiers (Supplementary Table 1).

When cell lines are treated with a concentration range of Trametinib they reach a plateau phase in viability at higher concentrations (Supplementary Figure 1), suggesting that the pathway is completely inhibited here. Since no toxicity was observed a concentration of 1uM was chosen for the screen. Cell lines were screened with 10, 100 nM and 1 uM of the library compounds with and without Trametinib. All cell lines showed significant growth inhibition upon treatment with 1uM Trametinib (Supplementary Figure 2). To identify compounds that are more effective when combined, viability was normalized to solvent treated cells in the DMSO assays and to the Trametinib treated cells in the Trametinib assays. The Area Under the Curve (AUC) for all compounds was determined in both assays and subsequently normalized to the AUC for DMSO treated cells (Figure 1A).

**Figure 1:**
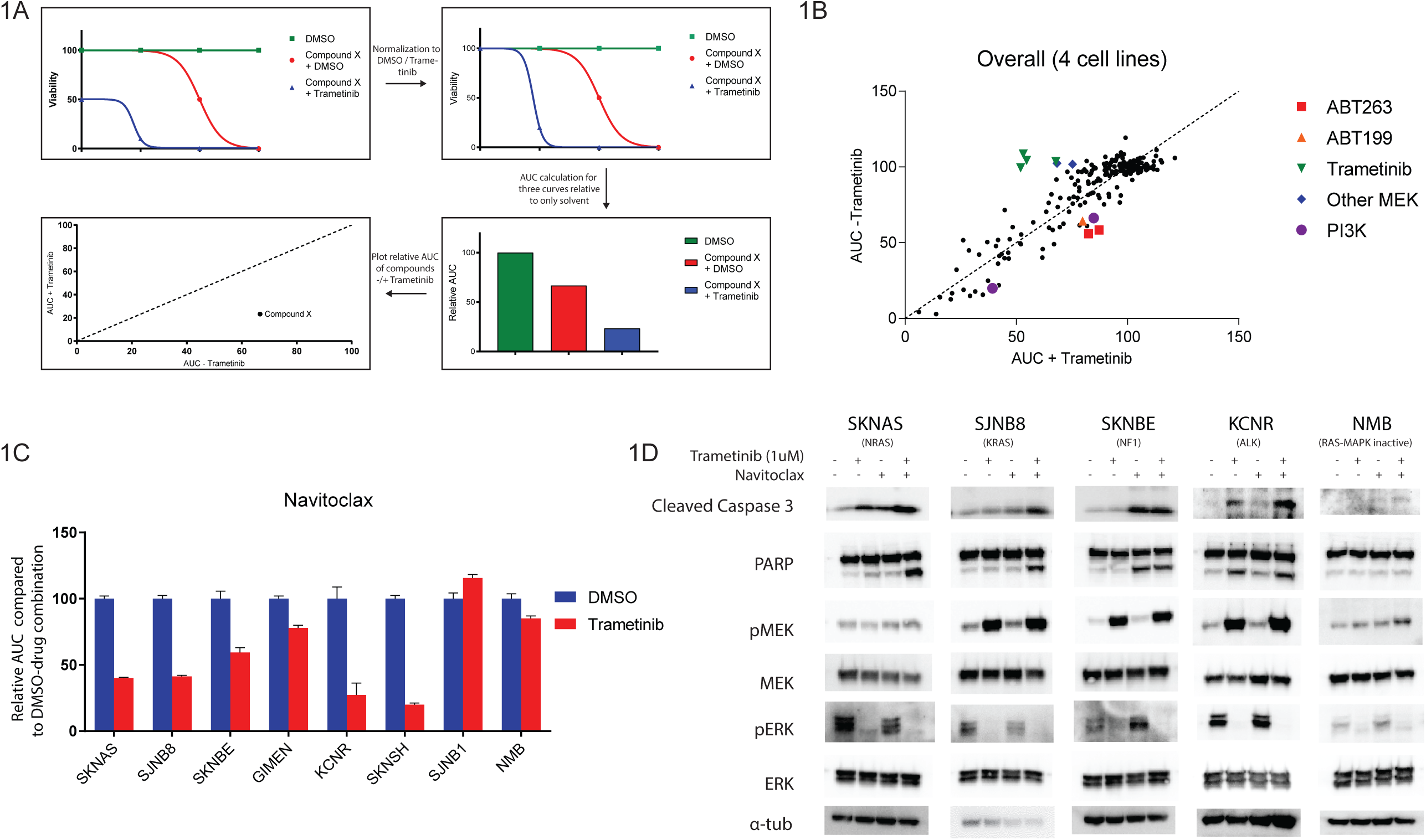
High throughput compound screening identifies ABT263 or Navitoclax as a combination target with Trametinib in neuroblastoma with RAS-MAPK mutations. A) Methodology for calculating the relative area under the curve, and the representations shown in 1B, from viability curves. B) Plots showing relative area under the curve for each screened compound in the presence of solvent or Trametinib averaged over the four neuroblastoma cell C) Area under the curve for an extended panel of cell lines treated with 7 concentrations of Navitoclax combined with solvent or Trametinib. The AUC was normalized to the AUC of cells treated with solvent and Navitoclax. D) Western blots of cell lines treated with Trametinib, Navitoclax or the combination for 48 hours. Cells treated with the combination show increased PARP cleavage and an increase in Cleaved Caspase 3.

Most compounds are similarly effective in the DMSO and Trametinib treated cells as evidenced by an average slope of 1.0 (range 0.95 tot 1.13) calculated by linear regression over all points per cell line (data not shown). As a negative control, we determined the effect of combining Trametinib with other MEK inhibitors. In cells treated with Trametinib, increased inhibition of RAS-MAPK signaling by the MEK inhibitors present in the library showed no additional effect (Figure 1B, Supplementary Figure 3). Contrastingly, PI3K inhibitors are known to work synergistically with MEK inhibitors^14^ and indeed have a larger effect when combined with Trametinib. This shows our screening approach is able to detect synergistic and antagonistic effects.

### Trametinib treatment sensitizes RAS-MAPK mutated neuroblastoma cells to Navitoclax

To rank the compounds in their effectiveness we calculated the average difference in AUC in the DMSO and Trametinib assay over the four cell lines (Supplementary Table 2). The most effective compound was ABT263 or Navitoclax. Navitoclax is an inhibitor of the interaction of Bim and the anti-apoptotic family of Bcl2 like proteins^15^. Its analog compound ABT199 or Venetoclax, which specifically targets Bcl2 and not the Bcl-XL and Bcl-W family members^16^, showed a similar effect in the SKNAS and KCNR cell lines (Supplementary Figure 3). Both compounds are very effective in neuroblastoma cell lines and in vivo models^17-19^. Furthermore, both compounds have shown to work synergistically with MEK inhibition in several other cancer types^20-22^.

To confirm the sensitizing effect observed in the screen, an extended range of concentrations was tested in a larger panel of cell lines. Trametinib is only effective in cell lines with RAS-MAPK mutations (Supplementary Figure 4). In these cell lines a clear sensitizing effect of Trametinib on Navitoclax sensitivity was observed, while in non-mutated cell lines this was less pronounced, suggesting that this effect is dependent on an active RAS-MAPK pathway (Figure 1C, Supplementary Figure 5).

Trametinib treatment of RAS-MAPK mutated cells has a cytostatic effect, while the combination with Navitoclax is expected to lead to increased apoptosis. In line with this, Trametinib treatment does not cause an increase in cleaved PARP nor cleaved Caspase 3 in most cell lines. Combining Trametinib with Navitoclax does cause an increase in PARP cleavage and cleaved Caspase 3 on Western blot, except in SKNBE, where the effect of Navitoclax alone is already quite strong (Figure 1D). The KCNR cell line is sensitive to Navitoclax, however, the combination with Trametinib still causes a clear increase in especially Caspase 3 cleavage. The NMB cell line, which does not contain RAS-MAPK mutations, does not respond to the combination.

### The sensitizing effect of Trametinib to Navitoclax is mediated by stabilization of Bim

Inhibition of the RAS-MAPK pathway is known to causes an increase in Bim protein levels, which is expected to lead to an increase in cell death. However, sequestration of Bim to anti-apoptotic proteins such as Bcl2, Bcl-XL and Mcl1, can prevent apoptosis, explaining the possible mechanism of synergy between Navitoclax and Trametinib^20^. In cell lines with RAS-MAPK mutations we indeed observe increased Bim protein levels when treated with Trametinib (Figure 2A). This was not observed in lines without mutations, indicating that an active pathway is a prerequisite for this effect.

**Figure 2:**
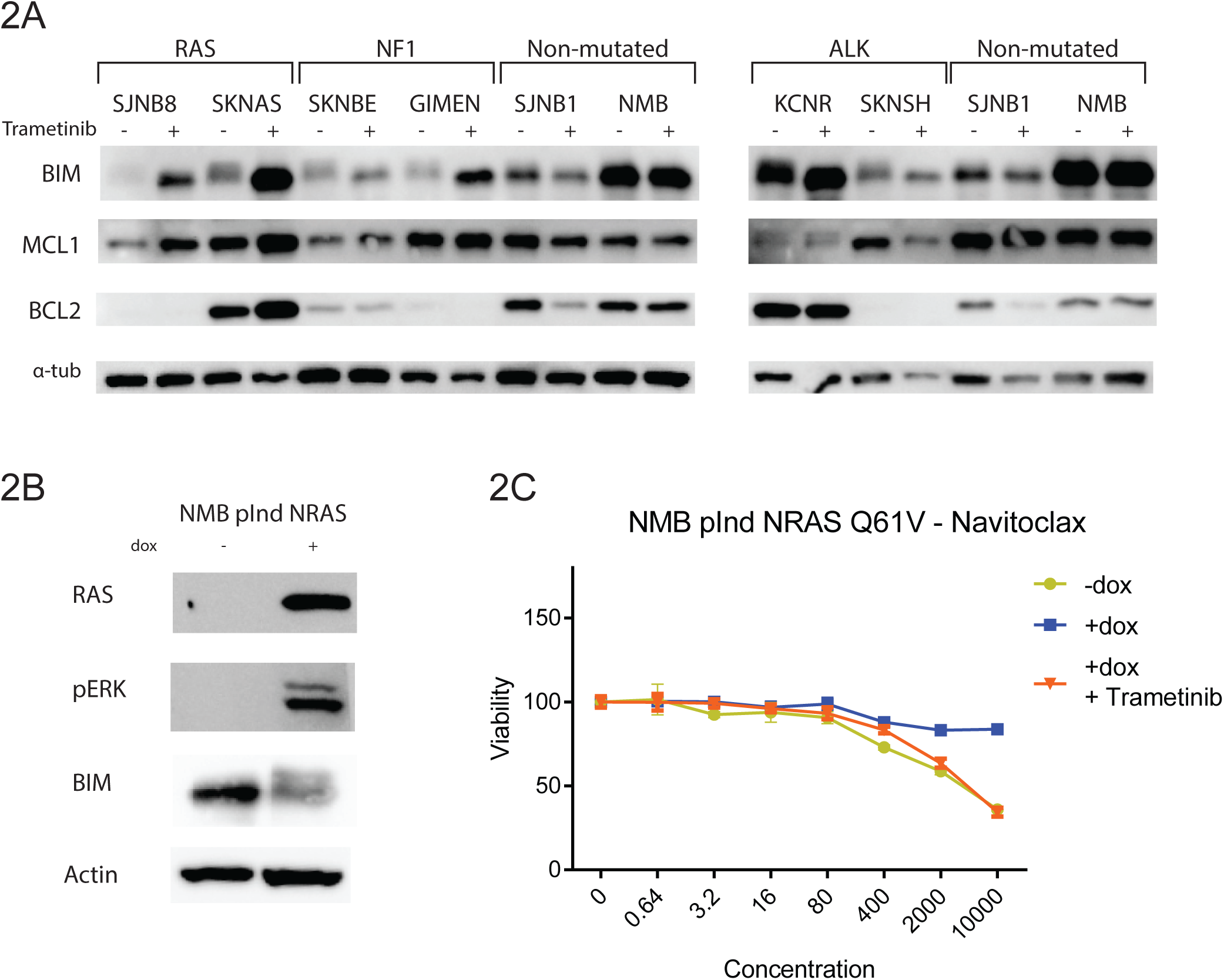
RAS-MAPK activation induces Bim destabilization and resistance to Navitoclax, which can be inhibited by MEK inhibitors. A) Western blot of cell lines treated with solvent or 1uM Trametinib for 6 hours. Cell lines are grouped based on their mutation status of RAS-MAPK associated genes. In lines with activating mutations MEK inhibition causes an increase in Bim levels, while this is not observed in cell lines without mutations. B) Western blot of inducible overexpression of NRAS Q61V in the cell line NMB. Expression causes activation of the RAS-MAPK pathway and a decrease in Bim. C) Cell viability curves of NMB lines with inducible overexpression of NRAS Q61V with and without Trametinib. Expression of mutant NRAS causes decreased sensitivity to Navitoclax and this sensitivity can be restored by Trametinib.

To test whether activation of the RAS-MAPK pathway has the opposite effect we expressed a mutant NRAS Q61V protein in the cell line NMB. This cell line does not have an active pathway and is insensitive to Trametinib. When the pathway is activated a decrease in total Bim levels is observed (Figure 2B), showing the involvement of this pathway. Expression of mutant NRAS also causes resistance to Navitoclax (Figure 2C), which can be abrogated by adding Trametinib to these cells, suggesting that it is indeed the MAPK pathway and not other pathways downstream of RAS that mediate this effect.

### Trametinib sensitizes neuroblastoma to various Bcl2 family member inhibitors

To further dissect specific effects of different BCL2 like proteins, S63845, a specific Mcl1 inhibitor, was also tested in our cell line panel. Most RAS-MAPK mutated lines were sensitized to S63845 by Trametinib (Figure 3A, Supplementary Figure 6). Furthermore, IP of Bim in SJNB8 also shows an increase in Mcl1 bound to Bim when treated with Trametinib (Supplementary Figure 7). This indicates that increased Bim binding to anti-apoptotic proteins could be a general mechanism for the sensitizing effect of MEK inhibitors to inhibitors of these proteins.

**Figure 3:**
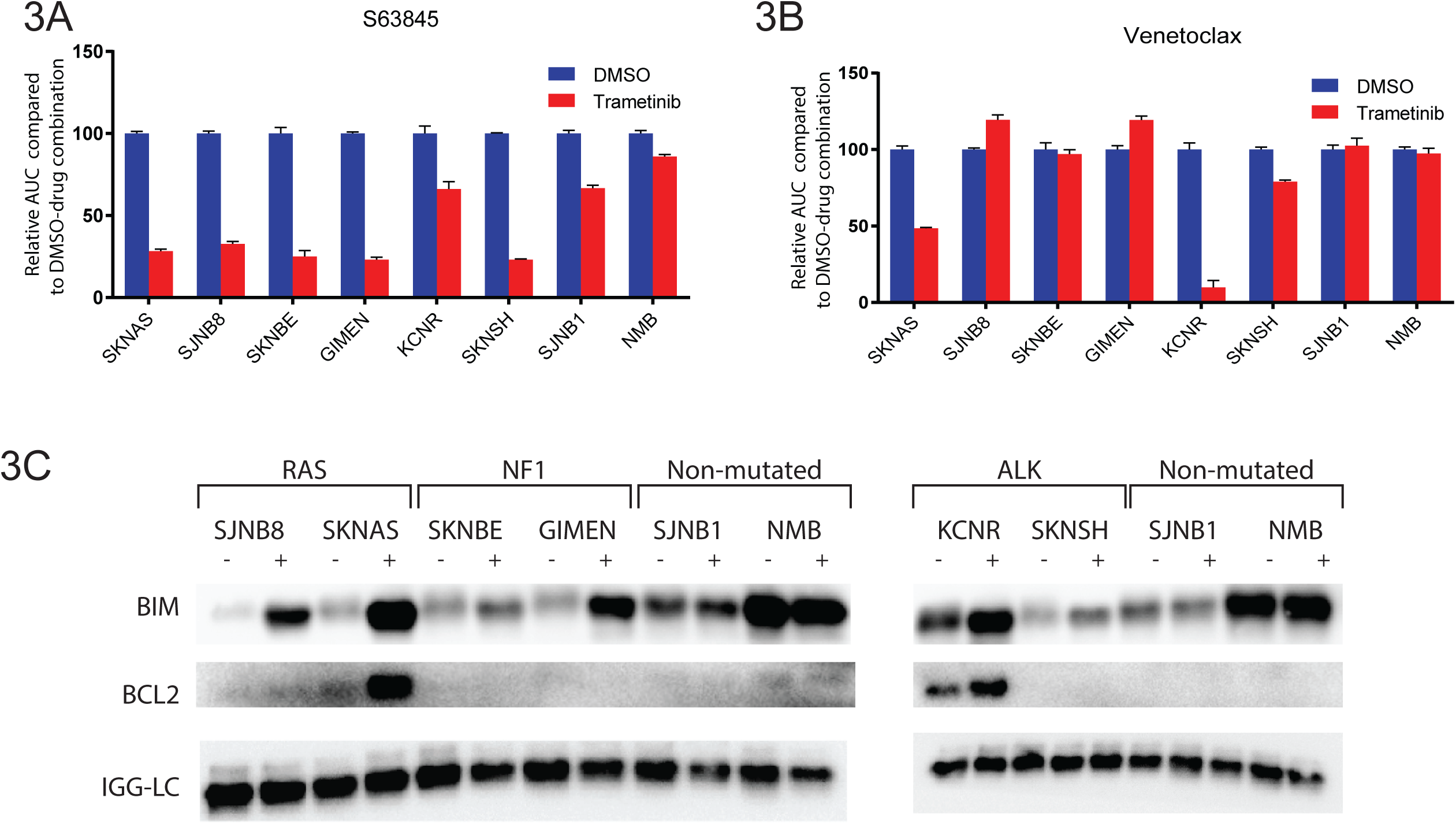
MEK inhibition sensitizes neuroblastoma cell lines with an active RAS-MAPK pathway to treatment with other Bcl2 family inhibitors. A) Area under the curve for a panel of cell lines treated with seven concentrations of the Mcl1 inhibitor S63845 combined with solvent or Trametinib. The AUC was normalized to the AUC of cells treated with solvent and S63845. B) Area under the curve for a panel of cell lines treated with seven concentrations of the specific Bcl2 inhibitor Venetoclax combined with solvent or Trametinib. The AUC was normalized to the AUC of cells treated with solvent and Venetoclax. SKNAS and KCNR show clear sensitization to Venetoclax by Trametinib. C) Western blot of Bim co-immunoprecipitation in lines treated with solvent or 1uM Trametinib for 6 hours. The KCNR cell line shows Bim bound to Bcl2 irrespective of the presence of Trametib, while the SKNAS cell lines shows Bim bound to Bcl2 only when Trametinib is present.

We also tested the sensitivity of the different cell lines to Venetoclax, the more specific Bcl2 inihibitor, in the absence and presence of Trametinib. (Figure 3B, Supplementary Figure 8). Only KCNR and SKNAS were sensitized toVenetoclax by Trametinib treatment. Co-immunoprecipitation experiments show BIM bound to Bcl2 only in these lines (Figure 3C), confirming that this binding can be used as a biomarker for Venetoclax sensitivity as was previously described^17^.

### Trametinib and Bcl2 family member inhibitors work synergistically in neuroblastoma xenografts

To validate the identified combination *in vivo*, we treated SKNAS xenografts with Trametinib and Navitoclax. Trametinib causes a significant difference in tumor volume compared to controls at day 7 and 21 (Figure 4A,B), which is expected since SKNAS has an *NRAS Q61V* mutation. Navitoclax alone had no significant effect on tumor growth, while tumors treated with Trametinib and Navitoclax were significantly smaller than tumors treated with Trametinib alone at the last time point (Figure 4A,B), showing that this combination also works in vivo.

**Figure 4:**
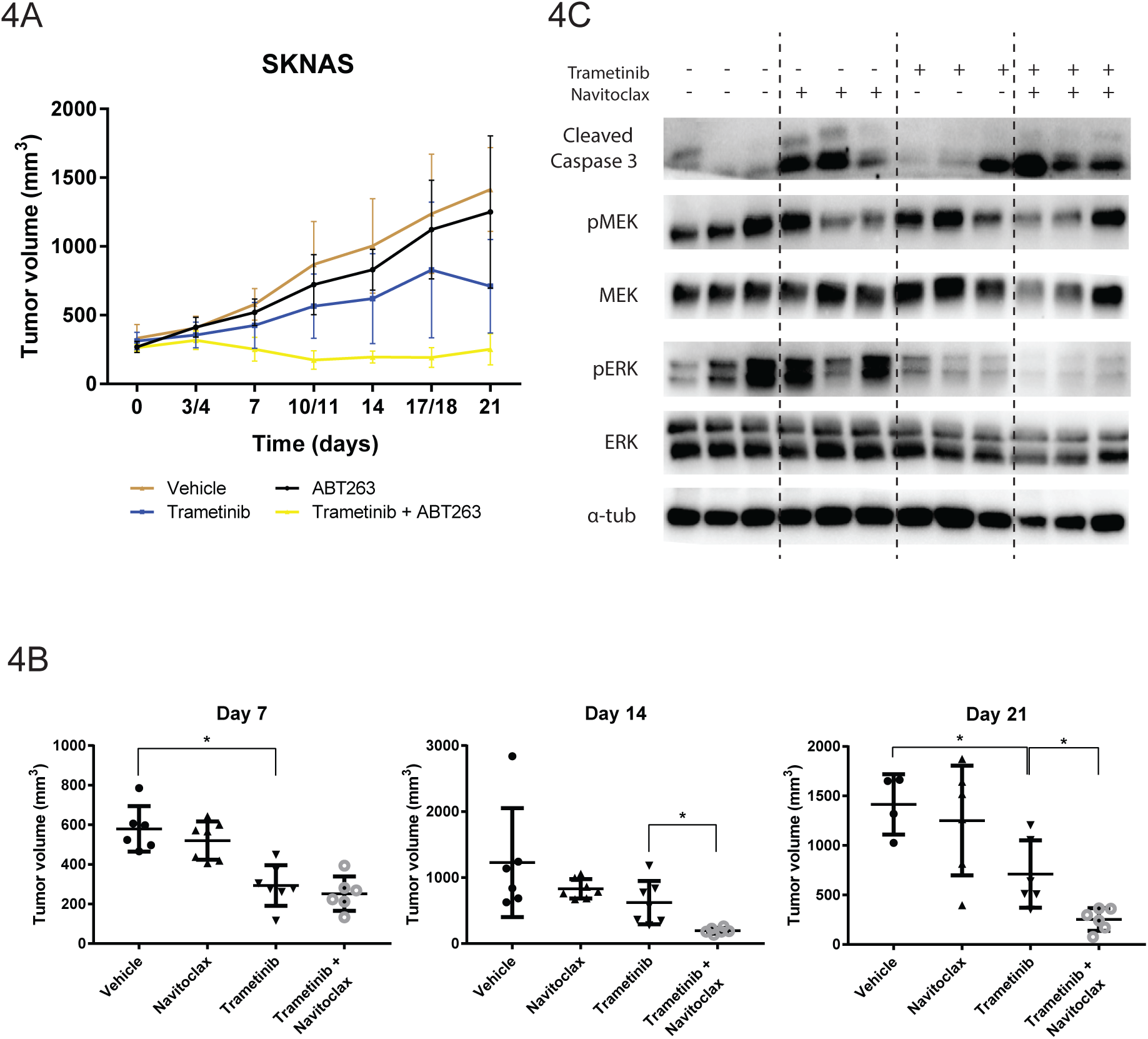
Navitoclax synergizes with Trametinib *in vivo* in SKNAS neuroblastoma xenografts A) Growth curves of xenografts treated with the indicated combinations. B) Tumor size at day 7, 14 and 21 by treatment group. P-values were calculated by Students t-test. * indicates p-value <0.05) C) Western blot of tumors from the different treatment groups. Treatment with Navitoclax causes an increase in Cleaved Caspase 3, while Trametinib inhibits the RAS-MAPK pathway as indicated by decreased pERK.

Western blot analysis shows that Navitoclax treatment causes an increase in cleaved Caspase 3, while this does not result in decreased tumor growth. Trametinib causes a decrease in ERK phosphorylation, which does result in decreased tumor growth, but not in an increase in cleaved Caspase 3. The combination causes a decrease in phosphorylated ERK and an increase in cleaved Caspase 3, although not significantly more than Nacvitoclax alone. This indicates that Trametinib works on target and that the combination causes increased apoptosis compared to Trametinib alone.

The same experimental set-up was used to test the combination of Trametinib and Venetoclax, since it is less toxic in clinical use. There was a trend that tumors treated with Venetoclax and Trametinib were smaller than the ones treated with Trametinib alone, although the difference was not significant (Supplementary Figure 9, p=0.07 at t=14). This shows that this combination could hold potential, although it is less powerful than Trametinib and Navitoclax.

## Discussion

We identify Navitoclax as a potential combination strategy with Trametinib in RAS-MAPK mutated neuroblastoma tumors which works through stabilization of Bim. This mechanism was previously reported in other tumors^20, 23^, and conversely, that RAS and *BRAF* mutations have destabilizing effects on Bim^24, 25^. Here we show that pathway activation through *NF1* deletion and *ALK* mutation also has a destabilizing effect, and that activation of this pathway leads to resistance to Navitoclax treatment. This shows that Bim destabilization by RAS-MAPK activation is a common mechanism and that a wide range of RAS active cancers could benefit from combination of MEK and Bcl2 family member inhibition.

Navitoclax has shown great efficacy in preclinical studies^26, 27^. However, in clinical trials it became apparent that inhibiting the binding between Bim and Bcl-Xl causes severe thrombocytopenia, limiting the therapeutic potential of this compound^28^. An alternative is Venetoclax, a drug that selectively inhibits binding of Bcl2 to Bim and therefore lacks the side-effects of Navitoclax^16^, which has also been shown to be active in neuroblastoma cell lines with high *Bcl2*^17^. Venetoclax was present in the screen performed, and showed synergy with Trametinib, but only in the cell lines that have high *Bcl2* expression.

We observe a heterogeneous response to the different targeted Bcl2 family inhibitors in our cell line panel. Previously it was shown that sensitivity to these inhibitors is correlated to the anti-apoptotic binding partner of Bim^29^. In primary neuroblastoma tumors a similar heterogeneity in Bim binding partners was observed^30^, suggesting that not all tumors will respond to Venetoclax and that different specific inhibitors should be used in conjunction with MEK inhibitors. Our results suggest that Mcl1 inhibitors could be effective in a subset of tumors with lower Bcl2 expression where Bim is bound to Mcl1. Therefore, their effectivity should be investigated *in vivo* in neuroblastoma.

Relapse neuroblastoma tumors frequently show RAS-MAPK mutations, and also in primary tumors this pathway is commonly activated. Since most tumors also express high levels of Bcl2, a significant part of neuroblastoma tumors could benefit from the combination of MEK inhibitors and Bcl2 inhibitors. Clinical trials that are currently being performed in neuroblastoma will establish whether these compounds are effective in RAS-MAPK mutated and high *Bcl2* expressing tumors respectively. We would propose to combine these compounds in a clinical trial, as long as both compound work on-target, even if results for single trials are modest.

## Acknowledgements

The research in this paper was supported by the Kinder Kankervrij Foundation (KiKa) under grant nr 189 and the European Union via the TRANSCAN2 project TORPEDO.

## Conflict of interest statement

H.N. Caron is employed by Roche.

## Notes

#### Summary of Updates

The author Lindy Vernooij was added to the manuscript file, where she was erroneously omitted before.

