## Supplementary Figures for "MEK inhibition causes Bim stabilization and sensitivity to Bcl2 family member inhibitors in RAS-MAPK mutated neuroblastoma"

### Supplementary Figure 1

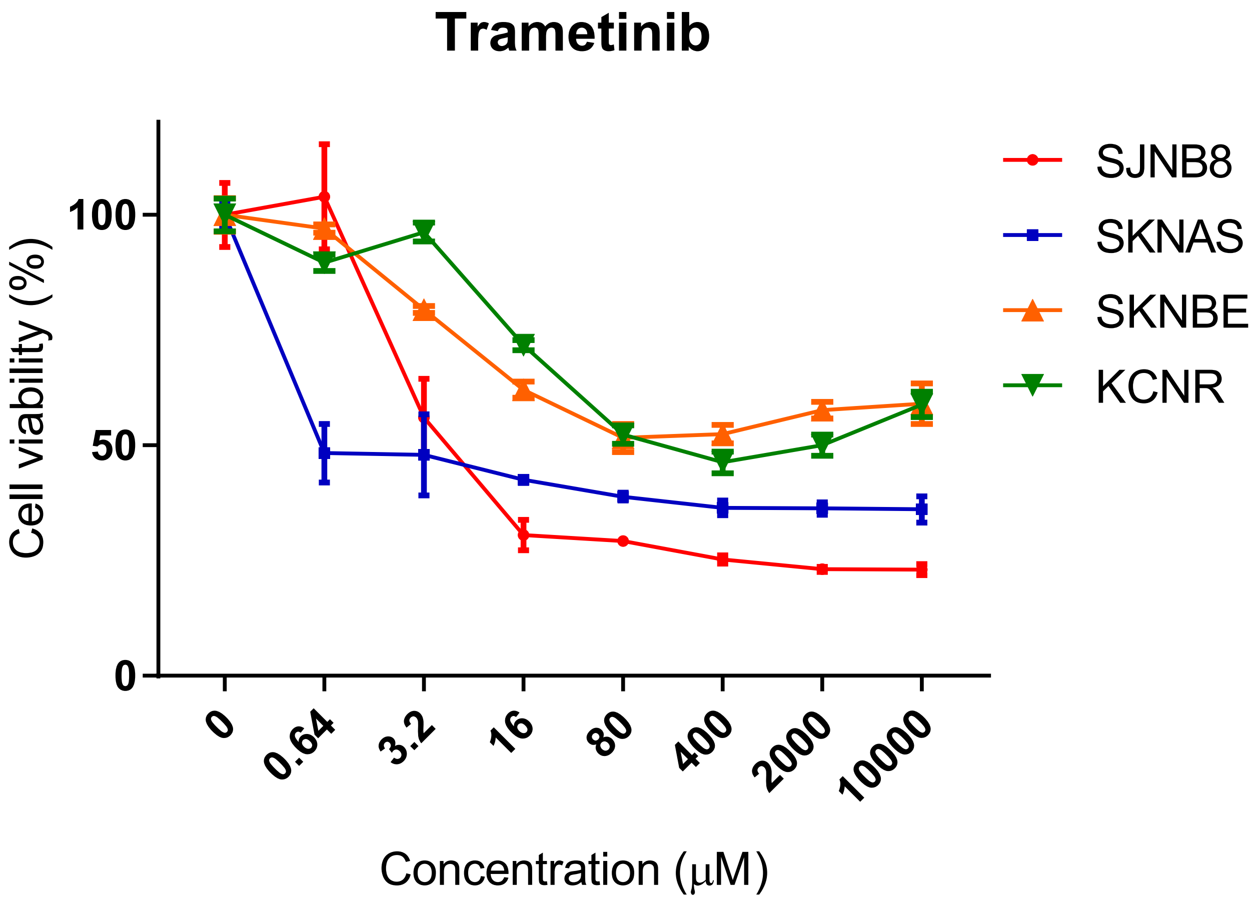


Supplementary Figure 1: Cell viability curves of the cell lines used in the screen treated with a concentration range of Trametinib.

### Supplementary Figure 2

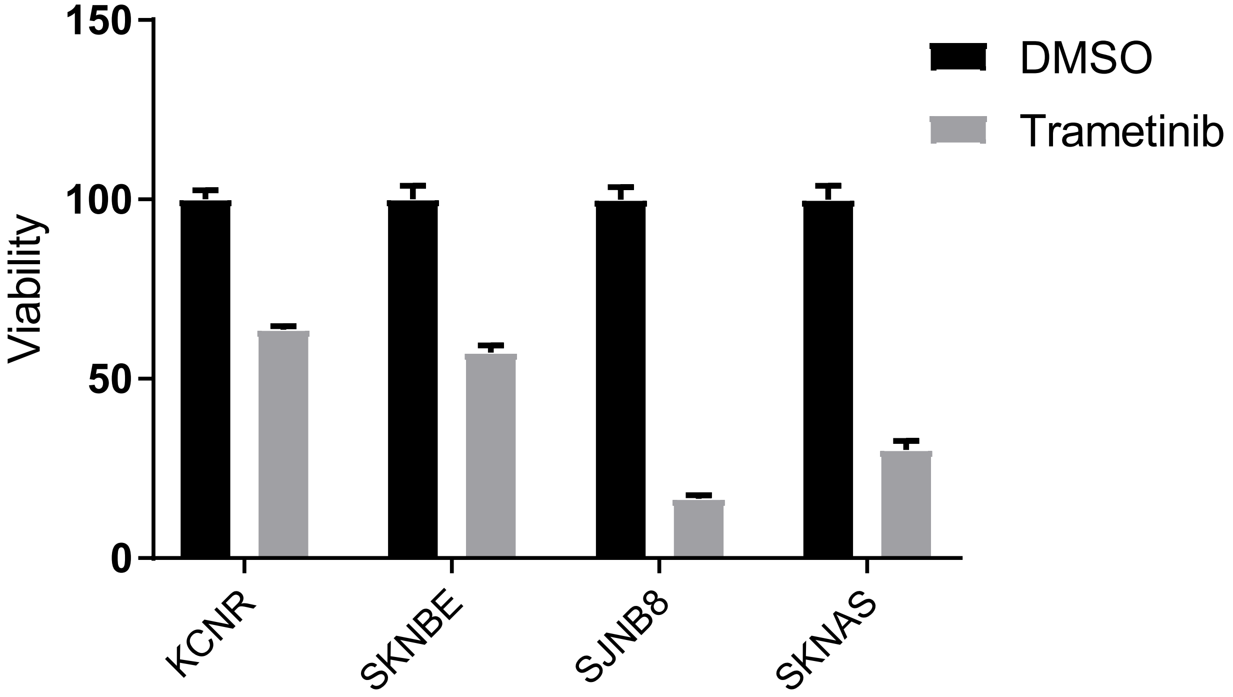


Supplementary Figure 2: Viability of the cells in the screen treated with 1uM of Trametinib or equivalent amounts of DMSO. Both conditions are normalized to the DMSO treated cells.

### Supplementary Figure 3

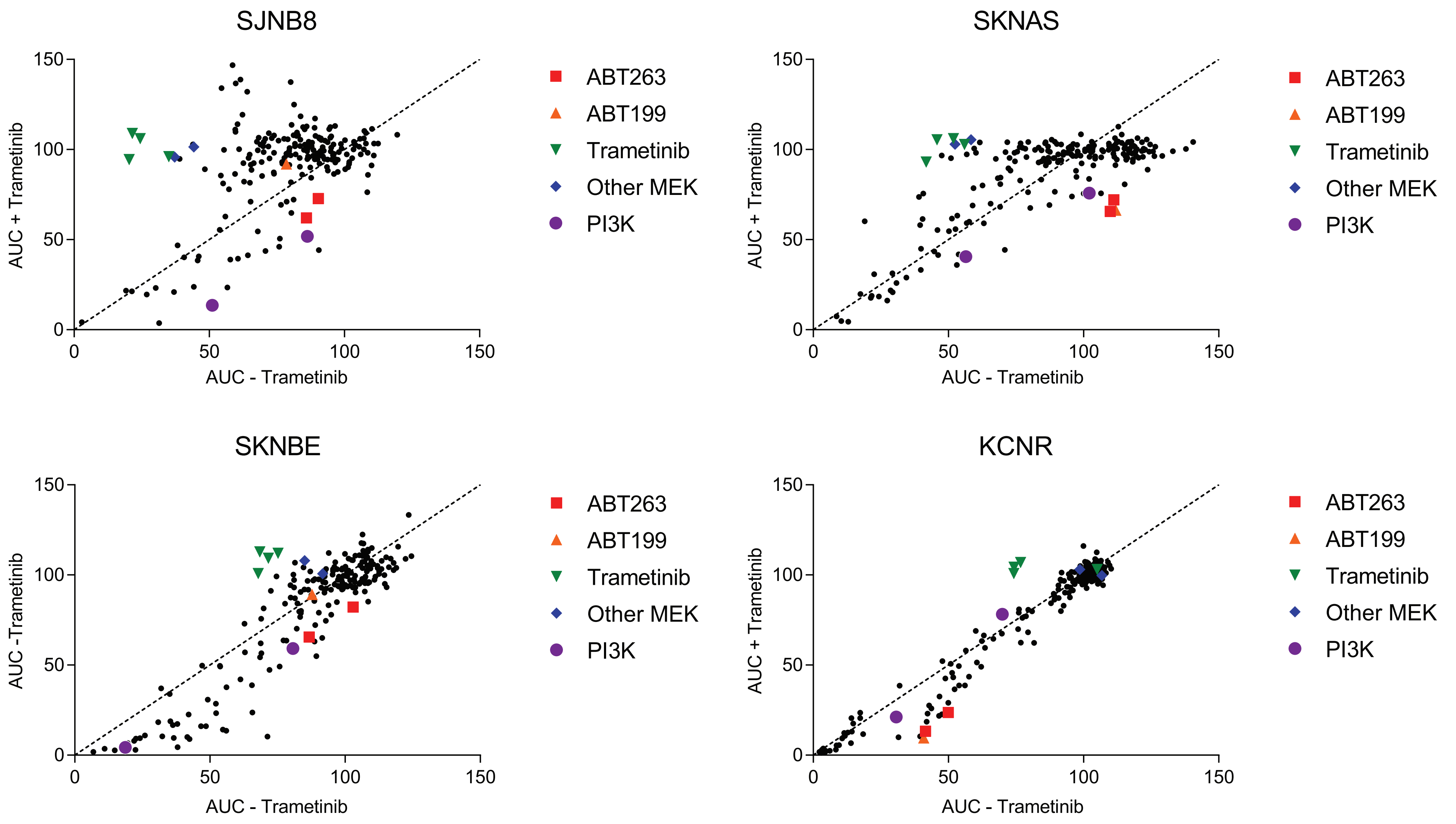


Supplementary Figure 3: Similar to figure 1B. Plots showing relative area under the curve for each screened compound in the presence of solvent or Trametinib in all four neuroblastoma cell line. Values were calculated as shown in figure 1A.

### Supplementary Figure 4

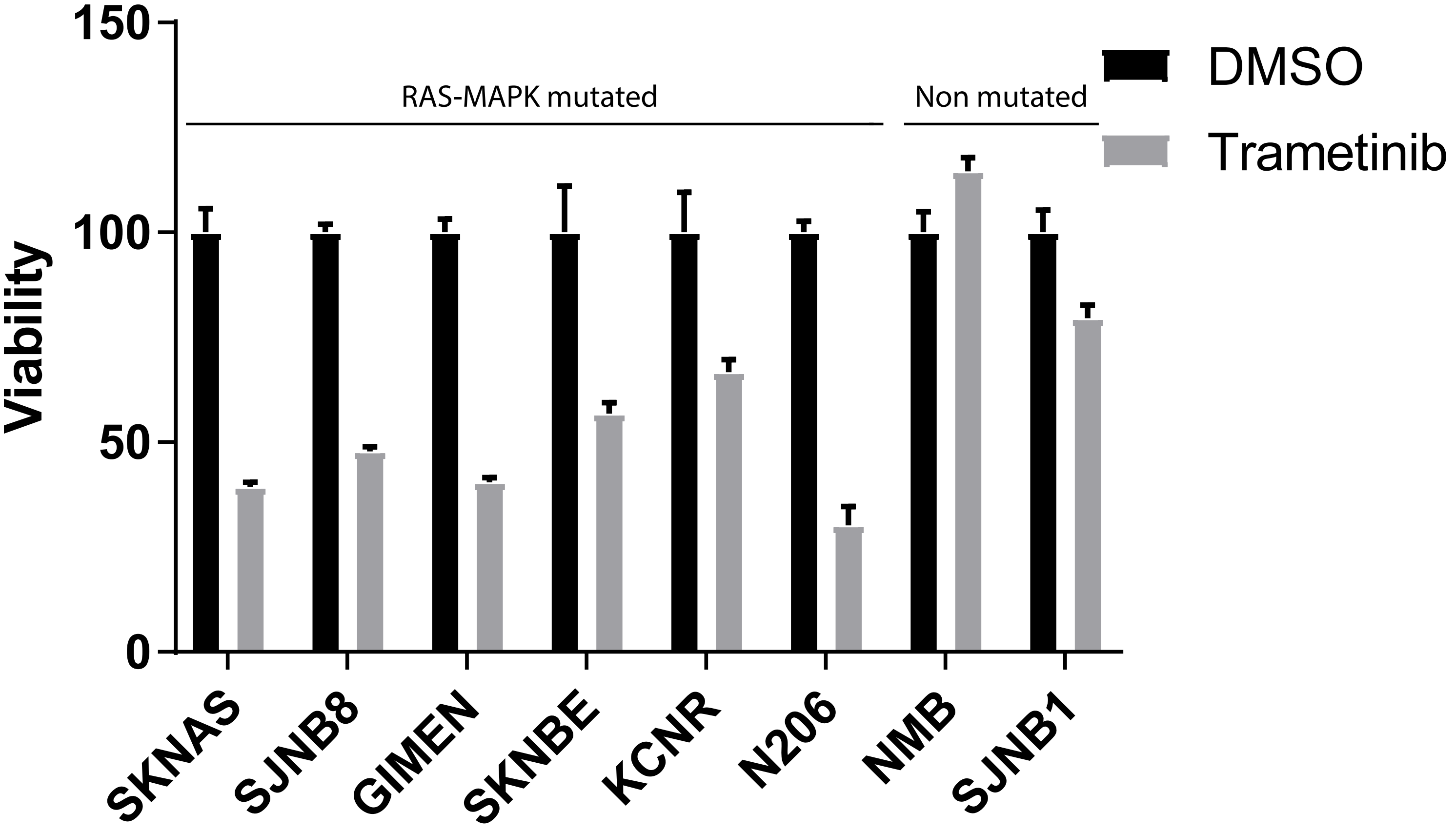


Supplementary Figure 4: Viability of the cell lines in the extended cell line panel treated with 1 uM of Trametinib or equivalent amounts of DMSO. Both conditions are normalized to DMSO treated cells.

### Supplementary Figure 5

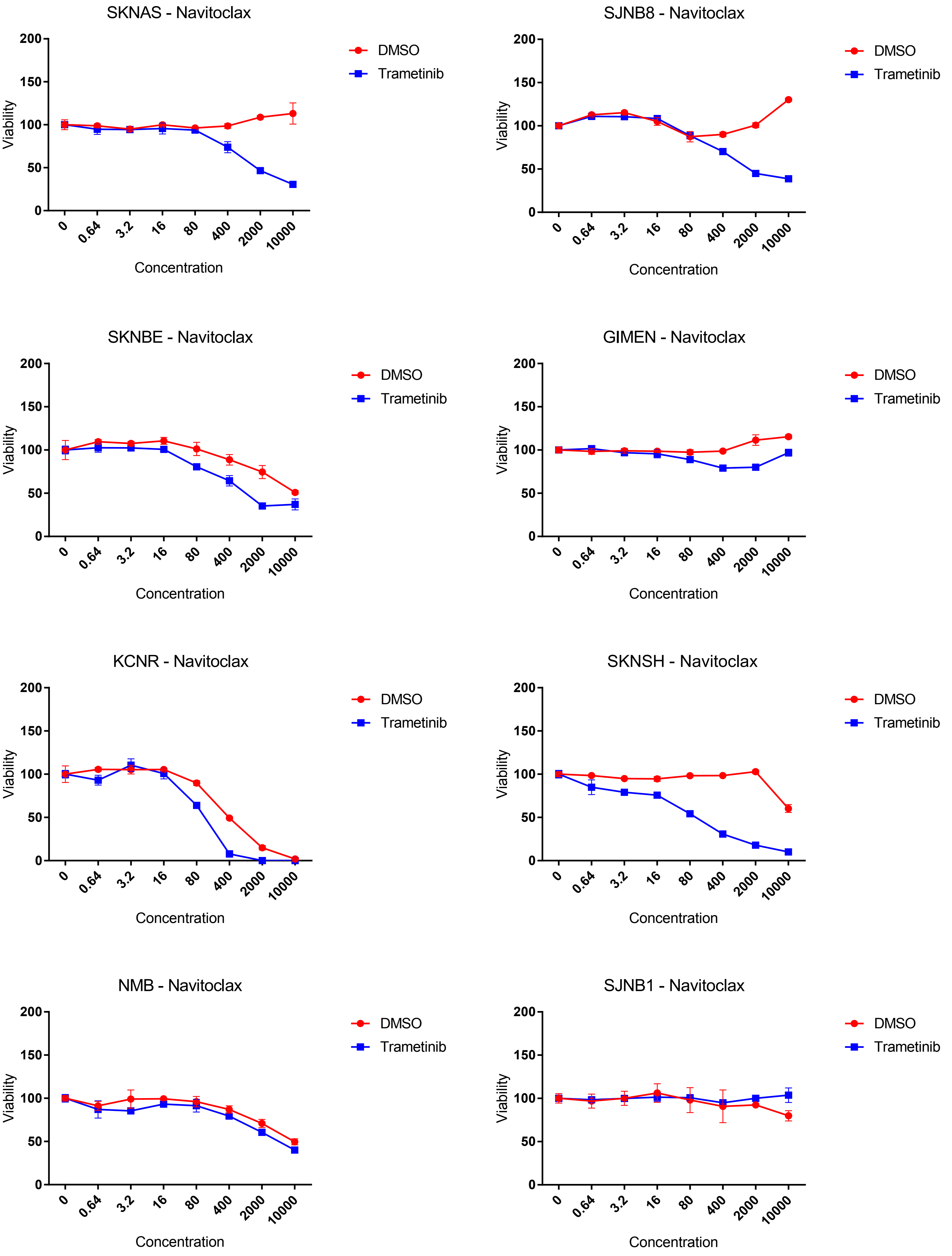


Supplementary Figure 5: Cell viability curves for Navitoclax in the presence and absence of 1 uM Trametinib. Values were normalized to Trametinib and DMSO treated cells respectively.

### Supplementary Figure 6

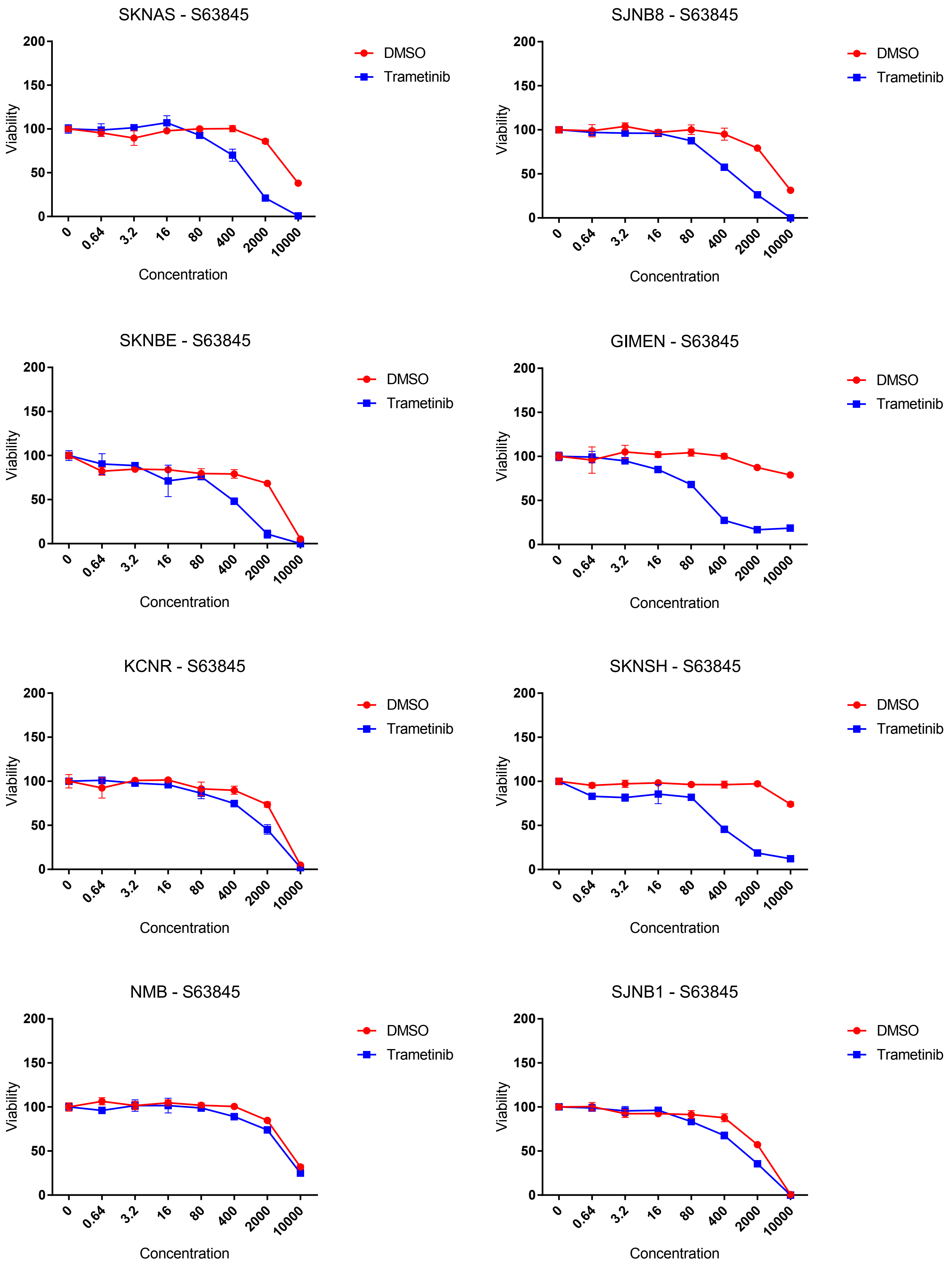


Supplementary Figure 6: Cell viability curves for S63845 in the presence and absence of 1 uM Trametinib. Values were normalized to Trametinib and DMSO treated cells respectively.

### Supplementary Figure 7

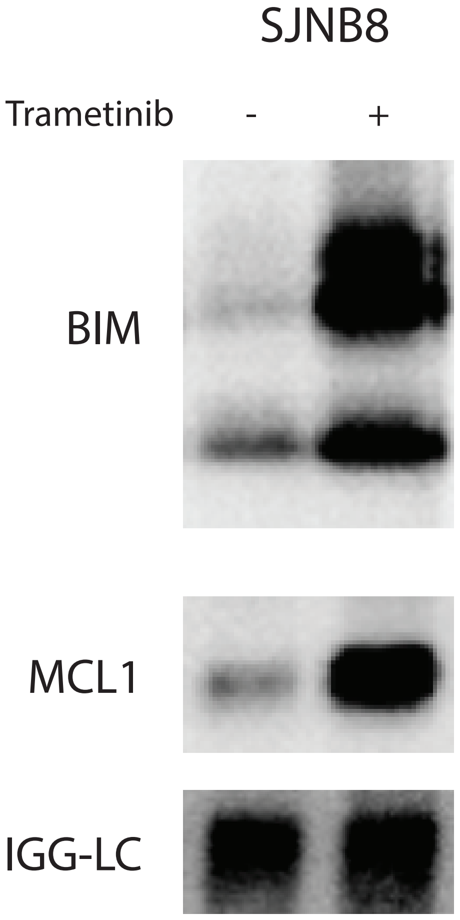


Supplementary Figure 7: Western blot of co-immuno precipitation of Bim in SJNB8 treated with Trametinib or equivalent amounts of DMSO.

### Supplementary Figure 8

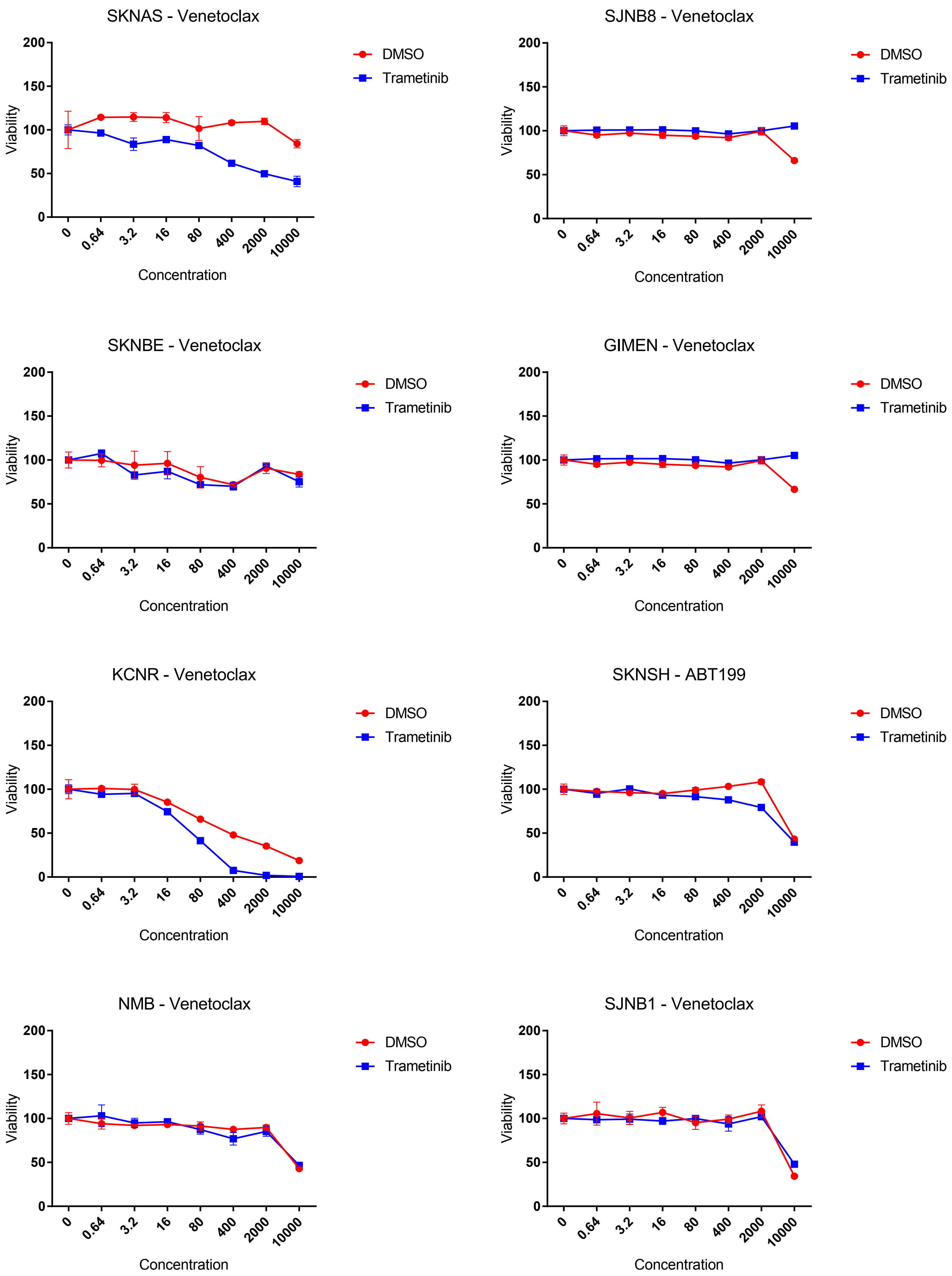


Supplementary Figure 8: Cell viability curves for Venetoclax in the presence and absence of 1 uM Trametinib. Values were normalized to Trametinib and DMSO treated cells respectively.

### Supplementary Figure 9

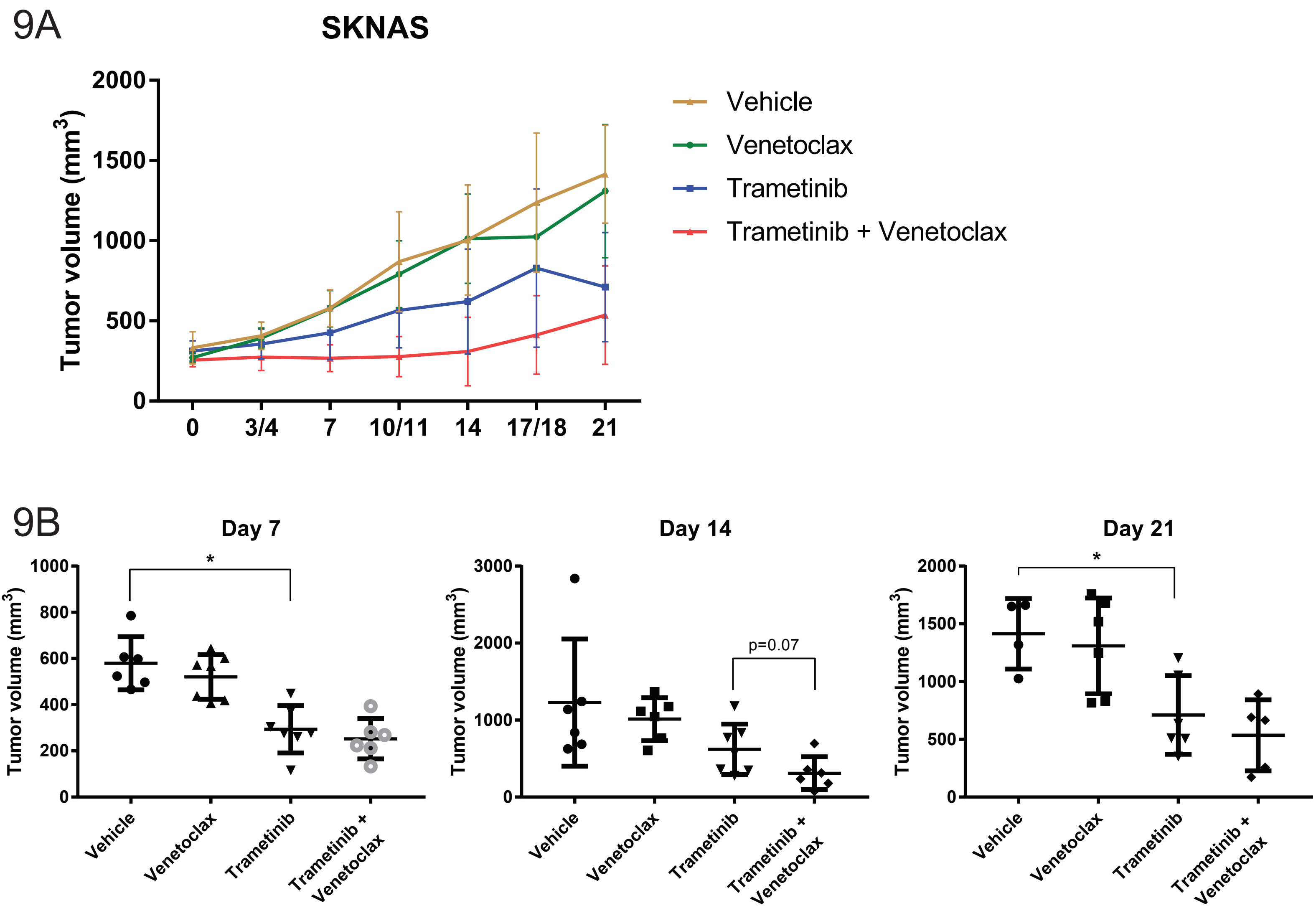


Supplementary Figure 9 In vivo testing of the combination of Trametinib and Venetoclax. Similar to Figure 4A,B. A) Growth curves of xenografts treated with the indicated combinations. B) Tumor size at day 7, 14 and 21 by treatment group. P-values were calculated by Students t-test. * indicates p-value <0.05)
